# Mesenchymal Stem Cells Sense the Toughness of Nanomaterials and Interfaces

**DOI:** 10.1101/2022.03.31.485540

**Authors:** Lihui Peng, Carlos Matellan, Armando del Rio Hernandez, Julien E. Gautrot

## Abstract

Stem cells are known to sense and respond to a broad range of physical stimuli arising from their extra-cellular environment. In particular, the role of the mechanical properties (Youngs or shear modulus, viscoelasticity) of biomaterials has extensively been shown to have a significant impact on the adhesion, spreading, expansion and differentiation of stem cells. In turn, cells exert forces on their environment that can lead to striking changes in shape, size and contraction of associated tissues, and may result in mechanical disruption and functional failure. However, no study has so far correlated stem cell phenotype and biomaterials toughness. Indeed, disentangling toughness-mediated cell response from other mechanosensing processes has remained elusive as it is particularly challenging to uncouple Youngs’ or shear moduli from toughness, within a range relevant to cell-generated forces. In this report, we show how the design of macromolecular architecture of polymer nanosheets regulates interfacial toughness, independently to interfacial shear storage modulus, and how this, in turn, controls the expansion of mesenchymal stem cells at liquid interfaces.

## Introduction

The mechanical properties of biomaterials have a significant impact on a wide range of cell phenotypes, from the regulation of cell spreading and cell proliferation to the modulation of fate decision^1-3^. In addition to cell response to the stiffness of their extra cellular environment, cells sense other mechanical features of their matrix, such as viscoelasticity^4-6^ and anisotropy^7,8^. These mechanical properties combine to ligand density^9^, nanoscale deformation^10^, matrix remodelling^11^ and other biochemical cues to trigger and regulate mechanosensing pathways^2,12^. Such sensing involves molecular force sensors^13,14^, directly enabling the probing of nanoscale mechanical properties of the cell microenvironments^15^. For example, cells have been found to respond directly to the local ligand density^16,17^ and to rearrange their local microenvironment, resulting in the regulation of cell spreading and tissue or organoid development^11,18^. In turn, mechanosensing processes, combined to cell contractility, regulate tissue formation, remodelling and function^19,20^. Poor control of these parameters may result in fracture and failure of biomaterials and interfaces, and associated impact on tissue repair^21-23^. However, little is known of the direct impact of materials toughness on cell phenotype, owing to the difficulty to uncouple toughness from other mechanical and physical parameters, at the cell scale and in a range relevant to cell-mediated contractile forces.

The importance of local mechanical properties of materials is clearly illustrated by the ability of cells to adhere, spread and proliferate at the surface of low viscosity liquids^24-28^. Indeed, it was demonstrated that fibroblasts, epithelial cells such as HaCaTs and keratinocytes, and mesenchymal stromal cells can proliferate at the surface of fluorinated oils, providing a mechanically strong protein nanosheet has formed at corresponding liquid-liquid interfaces. This enabled the formation of cell colonies at the surface of low viscosity liquid substrates that were as spread and dense as those formed on rigid tissue culture plastic and sustained the preservation of stem cell phenotypes^29-31^, despite the ultra-weak bulk mechanical properties of underlying substrates.

The impact that polymer and protein self-assembly has on liquid-liquid interfacial properties, including surface tension and interfacial pressure and viscosity is well established^32,33^. However, the impact of chemical and structural parameters of corresponding macromolecules on interfacial mechanics, and in turn cell spreading and phenotype at liquid interfaces, remain poorly understood. In this respect, protein-stabilised interfaces have been shown to display a broad range of interfacial mechanics and fluidity, resulting in the regulation of emulsion stability and associated formulations^34,35^. However, interfaces enabling the control of nanoscale mechanics and bioactivity, including cell adhesiveness, remain elusive.

Poly(L-lysine) (PLL) is a polycationic polymer that has been widely used for the functionalisation of a broad range of interfaces, enabling the direct coupling of other macromolecules and bioactive moieties as well as adsorption of extra-cellular matrix proteins. It was found to result in the formation of particularly stiff nanosheets at liquid-liquid interfaces, when assembling in the presence of reactive co-surfactants such as pentafluorobenzoyl chloride (PFBC)^30^ and, in turn, maintain the preservation of stemness and long term expansion of mesenchymal stem cells (MSCs)^29^. However, little is known of the structural parameters governing the mechanics of corresponding polymer/co-surfactant assembly and their nanoscale mechanics. In this study, we examine the role of molecular weight on self-assembly and interfacial mechanical properties of PLL nanosheets. We show that the molecular weight of PLL regulates interfacial toughness, resulting in interfaces displaying toughnesses comparable to that of steel, and enabling to resist cell-mediated contractile forces, for the formation of large and dense colonies.

## Results and Discussion

PLL assembles into stiff nanosheets at liquid-liquid interfaces when combined to reactive co-surfactants such as PFBC (Figure 1A). To investigate the adsorption and interfacial mechanical properties of nanosheets assembled from PLL with different M_w_, we used interfacial rheology (a du Noüy ring positioned at the liquid-liquid interface, coupled to a DHR3 rheometer). We first examined the impact of M_w_ on the adsorption of PLL at a fluorinated oil (Novec 7500)-aqueous (PBS) interface (Figure 1B). After equilibration of the system, PLL with different M_w_ was injected and the evolution of the interfacial shear moduli was monitored as a function of time. After a rapid initial increase, interfacial storage moduli gradually levelled within a range of 0.5-2.5 N/m. The kinetics of adsorption was found to depend on the molecular weight of PLL (Figure 1C). To quantify associated kinetics, we applied a Langmuir first order model^36^, assuming that interfacial shear moduli reflected the surface coverage at corresponding liquid-liquid interfaces. Adsorption traces were fit to the resulting equation:

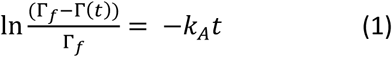

Where Γ(t) and Γ_f_ are the surface coverage of PLL at time t and equilibrium and k_A_ is the adsorption rate constant (see Methods). Our data was fitted over two separate early stages of the adsorption profiles, 100-600 s and 1500-2500 s (affording two rate constants, k_A1_ and k_A2_, respectively; Figure 1C-D). Although most traces fit a linear relationship, some deviation was clearly observed for the lowest molecular weight PLL tested (3 kDa). A gradual decrease in both rate constants was observed as a function of PLL M_w_, consistent with the expected impact of steric and coulombic hindrance associated with polyelectrolyte adsorption. However k_A1_ measured for 3 kDa PLL was significantly lower, presumably due to difficulty of achieving a percolated network at the liquid-liquid interface with low molecular weight molecules. In contrast, PLL chains with higher M_w_ can bridge across isolated adsorption islands more readily and may be expected to form a percolated network at early time points, following which stage the interfacial storage modulus may better reflect changes in polymer surface densities.

**Figure 1.**
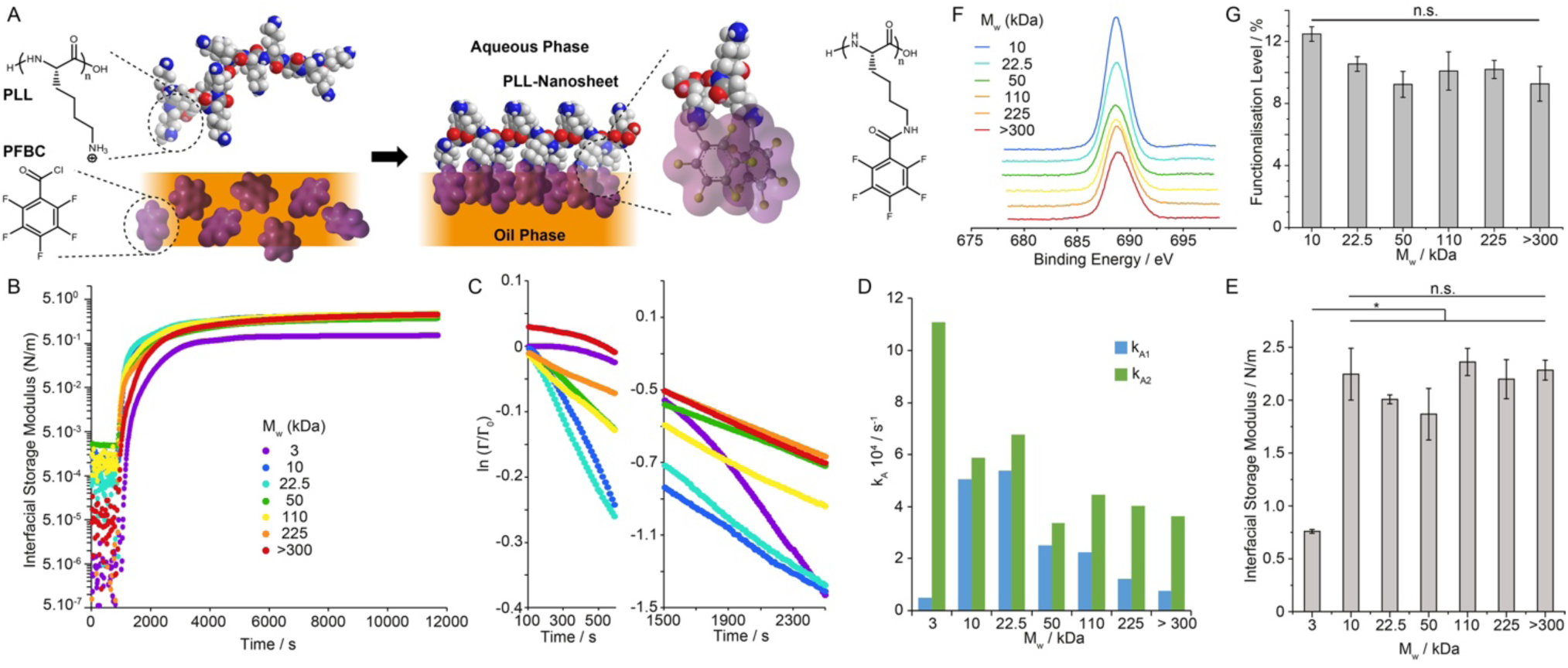
Impact of molecular weight on protein nanosheet assembly at liquid-liquid interfaces. A) Molecular structure of PLL nanosheets and proposed resulting architecture. B) Evolution of the interfacial shear storage modulus of PLL nanosheets forming at Novec 7500-water interfaces (Novec 7500 containing 10 μg/mL PFBC; aqueous solution is PBS with pH adjusted to 10.5; strain of 10^−3^ rad and 0.1 Hz). PLL with different M_w_ (3, 10, 22.5, 50, 110, 225 and >300 kDa) was introduced (after 900 s of equilibration) to make a final solution with a concentration of 100 μg/mL. C) Corresponding ln(Γ (t)/Γ _0_) plots at two different time points following protein injection. D) Adsorption rate constants extracted from corresponding linear fits. E) Interfacial storage moduli as a function of M_w_ of PLL, measured from frequency sweeps at a strain of 10^−3^ rad and 0.1 Hz. Error bars are s.e.m.; n≥3. F) XPS spectra (F 1s) obtained for nanosheets generated with PLL with different M_w_. G) Functionalisation levels quantified from corresponding XPS data (error bars are s.e.m.; n=3).

Despite differences in kinetics of adsorption, the ultimate (equilibrium) interfacial storage modulus of PLL interfaces was strikingly similar at different M_w_ (Figure 1E). The only interfaces displaying slightly lower interfacial shear storage moduli were those formed with 3 kDa PLL (0.76 N/m, compared to 2.0-2.3 N/m for higher M_w_). To examine whether assembled nanosheets were associated with changes in PLL adsorption densities, we characterised the abundance of PLL at corresponding interfaces, using tagged polymers and fluorescence microscopy (Supplementary Figure S1). This indicated comparable levels of polymer adsorption at liquid-liquid interfaces, independent of M_w_. Similarly, we characterised variations in the degree of functionalisation level of PFBC achieved on nanosheets from PLL with varying M_w_. To test this hypothesis, we characterised the atomic composition of PLL nanosheets by XPS (Figures 1F and G). Fluorination levels and associated PFBC functionalisation levels were found to be comparable for all nanosheet, independent of the molecular of the PLL used. The thickness of PLL nanosheets, characterised by neutron reflectometry in situ, was found to be in the range of 6-10 nm^37^. Therefore, we estimate the equivalent bulk shear modulus of materials that would be formed of PLL nanosheets to be in the range of 200-300 MPa. Such high stiffness implies the formation of a continuous rigid phase, which we propose is rich in rigid aromatic moieties able to aggregate via the formation of J-stacks (Figure 1a), owing to the strong quadripolar nature of fluorinated aromatics^38^. PLL nanosheets displaying comparable functionalisation with pentafluorobenzoate moieties would therefore be expected to display comparable interfacial storage moduli.

Considering the important role of viscoelasticity in the regulation of cell adhesion, migration and fate decision^6^, we next examined how the molecular weight of PLL impacted on the viscoelastic profile of nanosheets. Indeed, the interfacial storage modulus of PLL nanosheets displayed some frequency dependency associated with a clear viscoelastic response (Supplementary Figure S2). To characterise further viscoelasticity at PLL interfaces, we carried out interfacial stress relaxation experiments, using a double exponential decay model. Upon application of a defined strain (typically 0.1-1%), PLL nanosheets displayed apparent stress relaxation, with ultimate stress retention σ_R_ in the range of 55-75% (Figures 2A and B). Apart from nanosheets formed from 3 kDa PLL, our data indicated a gradual increase in the σ_R_, and therefore elasticity, as a function of increasing molecular weight.

**Figure 2.**
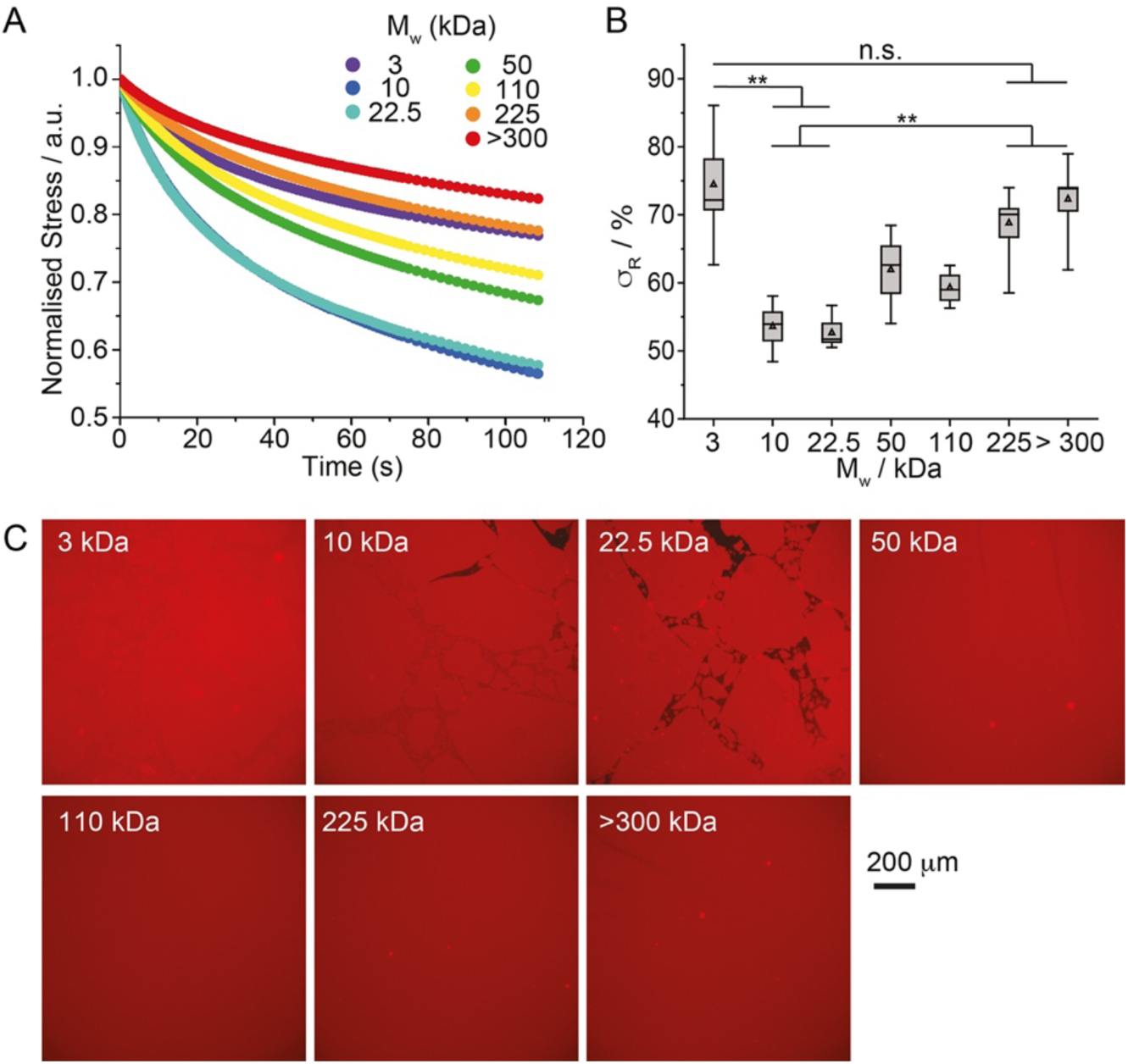
Interfacial viscoelasticity is controlled by the molecular weight of PLL. A) Representative stress relaxation profiles of nanosheets assembled from PLL with different molecular weights (strain: 10^−3^ rad). B) Corresponding stress retentions σ_R_ extracted from the corresponding fits. Error bars are s.e.m.; n⩾3. C) Epifluorescence microscopy images of PLL nanosheets assembled with PLL with a range of M_w_ (all tagged with Alexa Fluor 488). Detail of interfaces: Novec 7500 containing 10 μg/mL PFBC; aqueous solution is PBS with pH adjusted to 10.5; PLL with different M_w_ (3, 10, 22.5, 50, 110, 225 and >300 kDa) at a final concentration of 100 μg/mL.

This trend was surprising, considering the absence of change in interfacial storage modulus observed (Figure 1E) and, to gain further insight into this behaviour, we imaged corresponding liquid-liquid interfaces after formation of nanosheets assembled from tagged PLL with different M_w_ (Figure 2C). Although surface densities of PLL were found to be comparable (Supplementary Figure S1), the morphology of interfaces differed widely, dependent on PLL M_w_. Whereas at low molecular weight interfaces were apparently formed of fragmented nanosheets, they appeared homogenous and continuous over very large distances at higher molecular weights, with domains exceeding several millimetres. The transition to such behaviour was in the range of 50 kDa. Interestingly, although domains were clearly visible for the lowest molecular weight PLL tested (3 kDa), they remained tightly packed and no gap between such domains could be observed by microscopy. This may explain the high σ_R_ measured for nanosheets formed with 3 kDa PLL, and suggests that the viscous behaviour observed in PLL nanosheets results from inter-domain relaxation.

We next carried out interfacial creep experiments, again using the 6-element Burger’s model to quantify associated data (Supplementary Figure S3A-B). At low interfacial stress (1 mN/m), our data indicated a classic viscoelastic response (Supplementary Figure S3C), with no significant change in the main shear modulus G_0_ as a function of PLL M_w_, although G_1_ and G_2_ did increase slightly for PLL with M_w_ > 300 kDa (Supplementary Figure S3D). As in the case of stress relaxation experiments, the viscous component was found to increase significantly at intermediate M_w_ (Supplementary Figure S3E). However, at higher applied stress (5 and 10 mN/m), failure was clearly observed, depending on M_w_: interfaces formed with 50 kDa PLL failed at 10 mN/m, whereas those formed with 3 kDa PLL failed already at 5 mN/m (Supplementary Figure S3C). Hence nanosheet fracture and relaxation seemed to be strikingly impacted by the molecular weight of PLL.

Interfacial oscillatory rheology in amplitude sweeps was next carried out. PLL nanosheets displayed broad linear regions at low oscillation amplitudes, whereas significant thinning and non-linearity was observed at higher amplitudes. Such phenomenon is typical of the viscoelastic profile of concentrated polymer solutions and soft physically crosslinked polymer networks^39,40^ and was previously reported for other liquid-liquid interfaces stabilised by protein surfactants^35^. The toughness apparent from these amplitude sweeps, characterised from corresponding strain-stress traces, varied markedly depending on the molecular weight of the PLL forming the nanosheet (Figure 3A). We extracted interfacial toughnesses from these measurements, confirming a threshold of 50 kDa above which nanosheets were significantly reinforced (Figure 3B), in agreement with the nanosheet morphologies observed by fluorescence microscopy. Analysis of the damping function *h*(γ) associated with such non-linear viscoelastic profiles confirmed this threshold, with a significant shift in the position of the amplitude at which decay of the function and damping are observed (Figure 3C). In concentrated polymer solutions and at liquid-liquid interfaces, the damping functions are typically observed to collapse on the same profile, overlapping with the Soskey-Winter model^41^. In contrast, the striking shift in the damping function observed for M_w_ above 50 kDa, with overlapping functions for Mw > 50 kDa, implies a different mechanism for strain-induced softening. Indeed, although parameters associated with the Soskey-Winter damping model are not formally linked to molecular architectures (e.g. molecular weight, degree of crosslinking), the damping function is typically considered to reflect the mechanism of disruption of entanglements and physical bonds forming a polymer solution or network^35,40^. Therefore, the shift in damping function observed, by one order of magnitude in oscillation strain, is proposed to reflect a switch in domain relaxation and remodelling, from inter-domain to intra-domain rearrangement.

**Figure 3.**
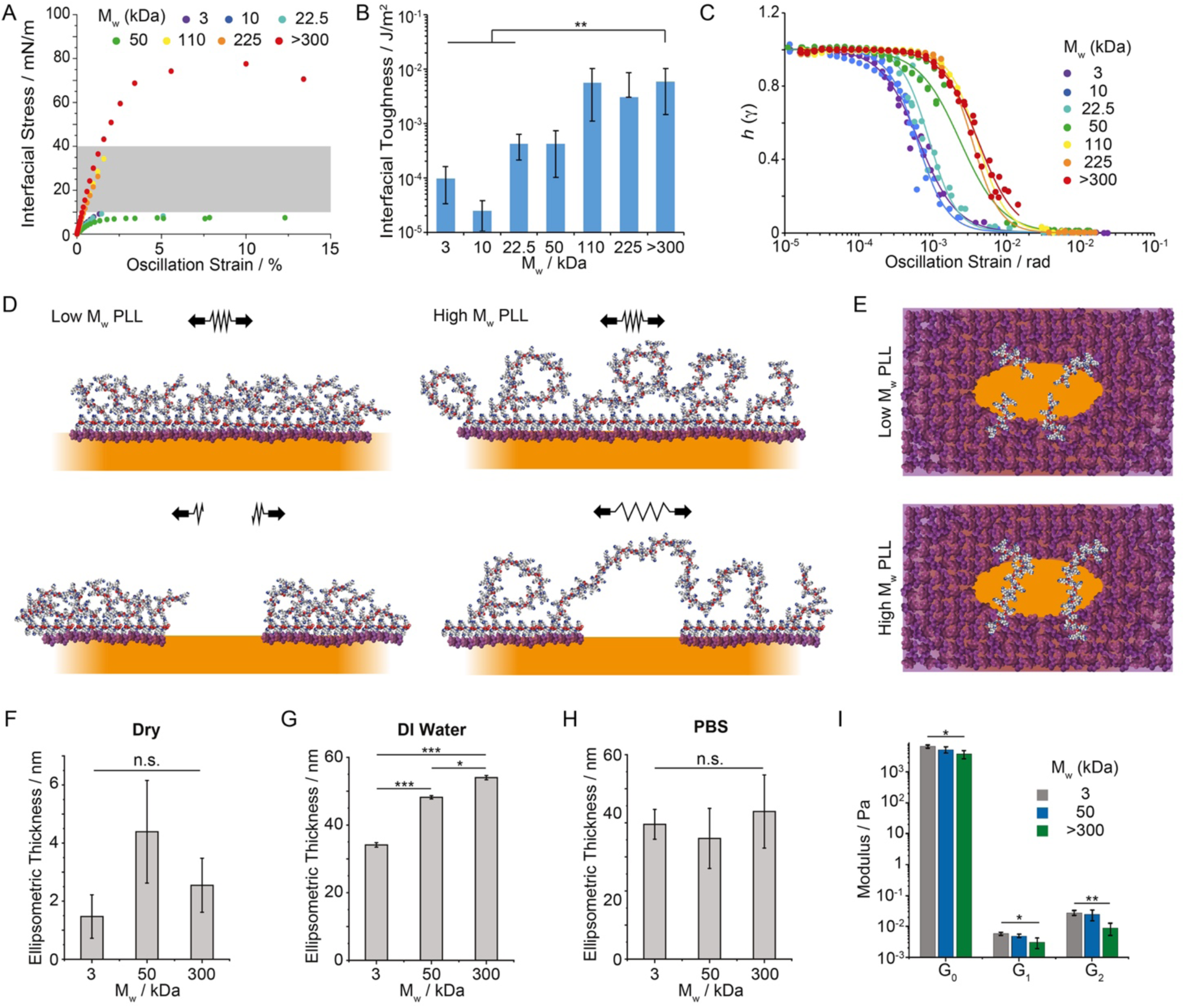
The nanoscale architecture of PLL nanosheets controls interfacial toughness. A) Representative shear stress-strain curves extracted from amplitude sweep experiments (frequency of 0.1 Hz). The grey area shaded correspond to the range of interfacial stresses expected to be exerted by mature focal adhesions. B) Summary of interfacial toughness calculated from the corresponding shear stress-strain profiles. (error bars are s.e.m.; n⩾3). C) Damping functions calculated from strain sweeps. The trend lines correspond to fits with the Soskey-Winter model. D and E) Proposed model of nanosheet fracture, depending on the molecular weight of PLL chains (D, side view; E, top view; only some chains localised at the fracture line are represented for improved visualisation). F-H) Ellipsometric thickness of selected nanosheets determined dry (F), in deionised (DI) water (G) and PBS (H). Nanosheets were transferred to silicon substrates using a Langmuir-Blodgett liquid-liquid trough, prior to characterisation. Error bars are s.e.m.; n = 3. I) Summary of magnetic-tweezer assisted interfacial microrheology data (shear moduli G_0_, G_1_ and G_2_ extracted using the 6 elements Burger’s model). Detail of interfaces: Novec 7500 containing 10 μg/mL PFBC; aqueous solution is PBS with pH adjusted to 10.5; PLL with different M_w_ (3, 10, 22.5, 50, 110, 225 and >300 kDa) at a final concentration of 100 μg/mL. Error bars are s.e.m.; n ≥ 7.

To further explore the mechanism of fracture mechanics and relaxation of PLL nanosheets with different M_w_, we characterised the thickness of nanosheets and their swelling. Nanosheets were transferred from corresponding liquid-liquid interfaces to mica and silicon substrates via Langmuir-Blodgett transfer. Tagged nanosheets were transferred to mica substrates in order to confirm via fluorescence imaging that large macroscopic nanosheets covering the surface of the target substrates could be transferred (Supplementary Figure S4). Nanosheets transferred to silicon substrates were then characterised by ellipsometry (Figure 3F-H). Dry nanosheets displayed thicknesses in the range of 1-4 nm. In deionised water, the hydrophilic phase of nanosheets increasingly swells, to 30-50 nm, depending on PLL M_w_. In contrast, in PBS, the swelling of all nanosheets was comparable and reduced compared to deionised water. Such ionic strength-dependent behaviour is expected from polyelectrolytes tethered to interfaces^42-44^ and increased hydrodynamic diameter associated with PLL chains of increasing M_w_^45,46^.

In addition, we characterised the nanoscale mechanical properties of the soft hydrophilic phase of PLL nanosheets via magnetic-tweezer assisted interfacial micro-rheology. Negatively charged magnetic particles were allowed to adhere to PLL nanosheets, prior to the application of a 30 s force pulse via a magnetic tweezer (Supplementary Figure S5). Bead trajectories were monitored (50 frames per second), the creep profile associated with such stimulation was then modelled using a 6-element Burger’s model and the associated shear moduli were quantified (Figure 3I). In contrast to the macroscopic interfacial rheology data obtained (Figure 1E and Supplementary Figure S3), the moduli of PLL interfaces were found to decrease as a function of molecular weight. Together with our ellipsometry data, this suggests that the hydrated, swollen soft phase of PLL nanosheets is increasingly soft and stretchable at high PLL M_w_. In turn, this soft phase is able to bridge across fracture cracks, dissipate local energy and reinforce the brittle PFBC-rich hard phase of PLL nanosheets (Figure 3D and E), in an analogous manner to polymer-reinforced composites^47-49^ and engineered tough hydrogels^50,51^.

We next examined how the toughness of PLL nanosheets may impact on stem cell adhesion and proliferation at liquid interfaces. Mesenchymal stem cells (MSCs) were allowed to adhere to fibronectin-coated PLL nanosheet-stabilised liquid-liquid interfaces and their spreading was characterised by immunostaining and confocal microscopy (Figure 4 A and B). Cells assembled a structured actin cytoskeleton on liquid substrates, despite the fluidity and low viscosity of the underlying substrate (Novec 7500). Morphological analysis revealed no significant difference between cells adhering to PLL nanosheets with different M_w_ and cells adhering to rigid glass coverslips coated with PLL and fibronectin. Similarly, MSCs formed focal adhesions located at the end of stress fibres on all PLL nanosheets, independent of the PLL M_w_ (Figure 4C).

**Figure 4.**
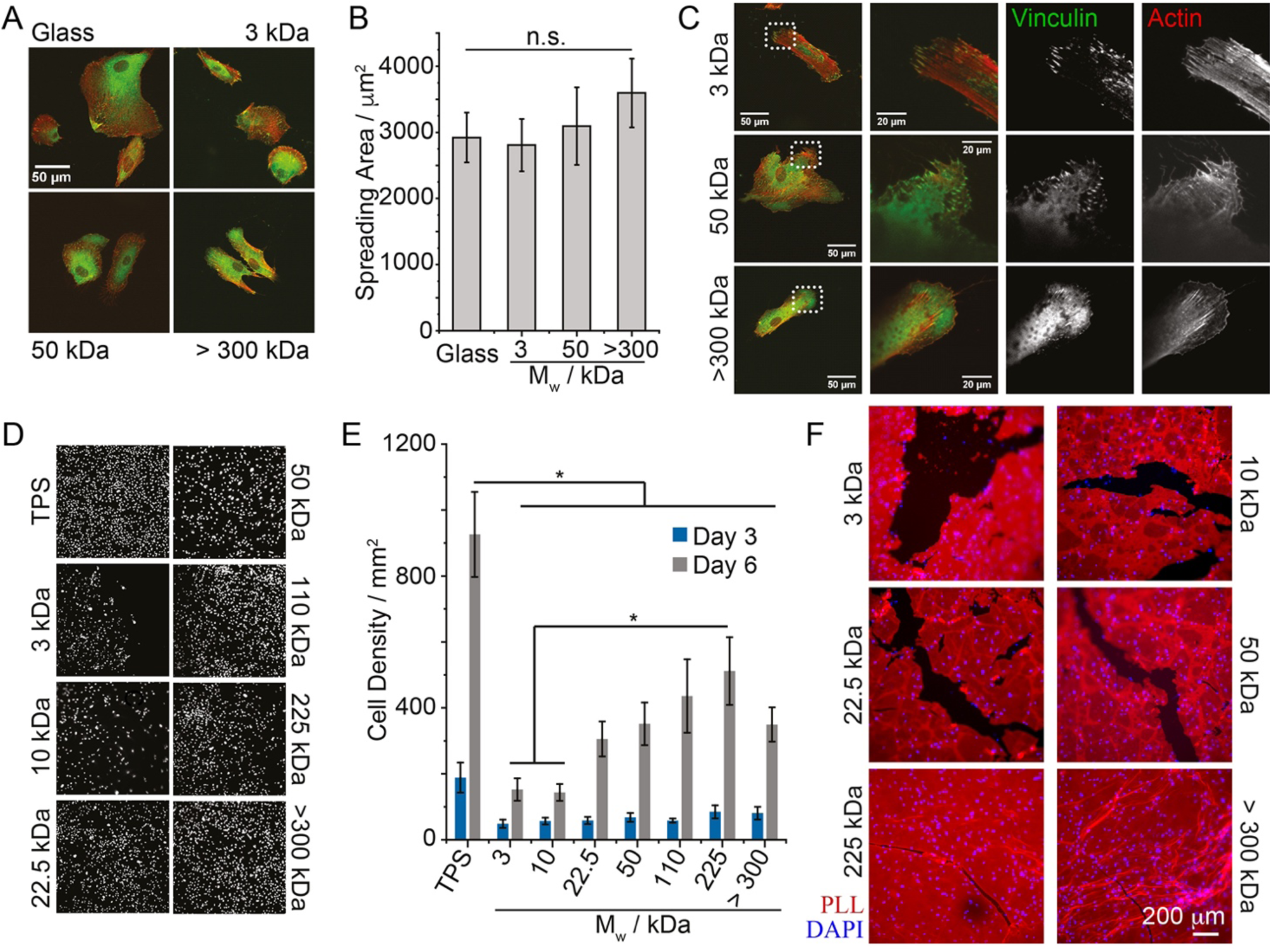
Stem cell expansion at liquid interfaces correlates with interfacial toughness. A and B) Impact of PLL molecular weight on cell spreading at Novec 7500 interfaces stabilised by corresponding nanosheets. C) Confocal microscopy images of MSCs spreading (after 24 h) on PLL/FN functionalised Novec 7500 interfaces. Zoom-in correspond to the dotted boxes. D and E) MSC expansion at PLL-stabilised Novec 7500 interfaces (D, representative nuclear stainings). F) Highly confluent MSCs remodel and fracture PLL/FN nanosheets assembled at the surface of Novec 7500. Epifluorescence microscopy images of PLL nanosheets 24 h after seeding MSCs at 200,000 cell/well (left). Red, PLL; blue, nuclei. Detail of interfaces: Novec 7500 containing 10 μg/mL PFBC; aqueous solution is PBS with pH adjusted to 10.5; PLL with different M_w_ (3, 10, 22.5, 50, 110, 225 and >300 kDa) at a final concentration of 100 μg/mL. Error bars are s.e.m.; n ≥ 4.

Therefore, consistent with the impact of hydrogels and biomaterials mechanics^52,53^, MSC adhesion to PLL nanosheet-stabilised interfaces with comparable interfacial storage moduli had no significant impact on cell adhesion and spreading. However, the proliferation of MSCs was significantly impacted by the molecular weight of PLL assembling nanosheets, and resulting interfacial toughness (Figure 4D ad E). This was not associated with any change in cell viability, which remained high and comparable on all substrates (Supplementary Figure S6). The abundance of fibronectin deposited at PLL nanosheet interfaces was also comparable (Supplementary Figure S7), in agreement with the similar spreading observed and the observation of focal adhesions and structured actin cytoskeleton on the different PLL interfaces tested. Instead, we propose that cells spreading on nanosheets assembled from lower M_w_ PLL sense the toughness of corresponding interfaces. Indeed, gaps in cell coverage were observed in cultures at nanosheets formed from low and intermediate M_w_ PLL. Hence, cell-mediated forces are proposed to locally fracture PLL nanosheets, or to extend shear induced fractures occurring during substrate preparation, leading to local relaxation of the 2D network and gradual reduction of the elasticity of the network.

Finally, to better evidence cell-mediated fracture of PLL nanosheets, we seeded MSCs at high densities at nanosheet-stabilised interfaces and characterised the morphology of resulting cultures 24 h after seeding. Clearer gaps can be seen in high density cultures seeded on low M_w_ PLL nanosheets (Figure 4F) and these gaps are clearly associated with fractures in PLL nanosheets. This suggests that local fracture in nanosheets, together with concerted contractile forces, further extending such defects to dimensions spanning several hundreds of microns, results in these gaps in dense cell cultures.

Overall, our data demonstrate that cells can directly sense the nanoscale toughness of interfaces to which they adhere and, despite developing mature adhesions at early time points, can mechanically disrupt their adhesive landscape, leading to retraction of adhesive areas and reduction in cell expansion. With forces exerted by cells in the range of 1-50 nN per adhesion, and focal adhesions displaying cross-section of 500 nm to 2 μm, the equivalent interfacial stress that can be expected to be transferred to nanosheets lies within the range of 5.10^−4^-0.1 N/m ^14,54-57^, and more specifically to the case of MSCs, fully mature adhesions were found to generate stresses near 10-40 mN/m (maximum forces in the range of 20-40 nN, with adhesions 1-2 μm in cross-section ^54,55^). This is precisely a range that is in between the ultimate interfacial stress that we measured for low and high M_w_ PLL nanosheets (Figure 3A). Therefore, the transition observed in interfacial toughness is proposed to overcome the maximum stress exerted by contractile cell adhesions, enabling to sustain nanosheet integrity over prolonged culture times.

The ability of cells to sense the mechanical properties of their environments is enabled by the reciprocal responses of the adhesion machinery (underpinned by integrin binding, actin assembly and contractility and mediated by adapter proteins such as talin and vinculin^13,58^) and the nanoscale mechanics of corresponding interfaces^9^. Deformation, strain stiffening and clustering, associated with the viscoelastic profiles of corresponding materials are integral elements sensed by cell adhesions and triggering downstream signalling. Our work proposes that the toughness of interfaces can further modulate such processes, not through the direct regulation of cell adhesion, but by defining a threshold above which interface and matrix remodelling lead to failure of the adhesive landscape, and as a result the retraction of cell adhered from associated areas. This concept is important to the engineering of biomaterials displaying significant mismatch in mechanical properties in bulk and at interfaces, as in the case of cell culture on liquid substrates and bioemulsions. It also underpins some of the processes occurring during matrix remodelling. For example, during the deposition of extra-cellular matrix and its mechanical integration to pre-existing biomaterials or tissues, or during tissue contraction in wound healing or tissue regeneration contexts.

## Supporting Information

Supporting Information is available from the author.

## Acknowledgements

Funding for this work from the European Research Council (ProLiCell, 772462), the China Scholarship Council (studentship, 201708060335) and the Leverhulme Trust Foundation for financial support (RPG-2017-229, Grant 69241) is gratefully acknowledged. We thank Dr Richard Whiteley for help with XPS.

## References

1 Guilak, F. et al. Control of stem cell fate by physical interactions with the extracellular matrix. Cell Stem Cell 5, 17–26 (2009).

2 Discher, D. E., Mooney, D. J. & Zandstra, P. W. Growth factors, matrices, and forces combine and control stem cells. Science 324, 1673–1677 (2009).

3 Trappmann, B. & Chen, C. S. How cells sense extracellular matrix stiffness: a material’s perspective. Curr. Opin. Biotech. 24, 948–953 (2013).

4 Chaudhuri, O. et al. Substrate stress relaxation regulates cell spreading. Nature communications 6, 6365 (2015).

5 Chaudhuri, O. et al. Hydrogels with tunable stress relaxation regulate stem cell fate and activity. Nat. Mater. 15, 326–334 (2016).

6 Chaudhuri, O., Cooper-White, J., Janmey, P. A., Mooney, D. J. & Shenoy, V. B. Effects of extracellular matrix viscoelasticity on cellular behaviour. Nature 584, 535–546 (2020).

7 Abhilash, A. S., Baker, B. M., Trappmann, B., Chen, C. S. & Shenoy, V. B. Remodeling of Fibrous Extracellular Matrices by Contractile Cells: Predictions from Discrete Fiber Network Simulations. Biophysical J 107, 1829–1840 (2014).

8 Mondrinos, M. J. et al. Surface-directed engineering of tissue anisotropy in microphysiological models of musculoskeletal tissue. Sci. Adv. 7, eabe9446 (2021).

9 Oria, R. et al. Force loading explains spatial sensing of ligands by cells. Nature 552, 219–224 (2017).

10 Attwood, S. J. et al. Adhesive ligand tether length affects the size and length of focal adhesions and influences cell spreading and attachment. Sci. Rep. 6, 34334 (2016).

11 Ferreira, S. A. et al. Bi-directional cell-pericellular matrix interactions direct stem cell fate. Nat. Commun. 9, 4049 (2018).

12 Dupont, S. et al. Role of YAP/TAZ in mechanotransduction. Nature 474, 179–183 (2011).

13 del Rio, A. et al. Stretching single talin rod molecules activates vinculin binding. Science 323, 638–641 (2009).

14 Pandey, P. et al. Cardiomyocytes Sense Matrix Rigidity through a Combination of Muscle and Non-muscle Myosin Contractions. Dev. Cell 44, 326–336 (2018).

15 Di Cio, S. & Gautrot, J. E. Cell sensing of physical properties at the nanoscale: mechanisms and control of cell adhesion and phenotype. Acta Biomater 30, 26–48 (2016).

16 Schvartzman, M. et al. Nanolithographic control of the spatial organization of cellular adhesion receptors at the single-molecule level. Nano Lett. 11, 1306–1312 (2011).

17 Deeg, J. A. et al. Impact of local versus global ligand density on cellular adhesion. Nano Lett. 11, 1469–1476 (2011).

18 Jowett, G. M. et al. ILC1 drive intestinal epithelial and matrix remodelling. Nat. Mater. 20, 250–259 (2020).

19 Rivron, N. C. et al. Tissue deformation spatially modulates VEGF signaling and angiogenesis. Proc. Natl. Acad. Sci. 109, 6886–6891 (2012).

20 Nawroth, J. C. et al. A tissue-engineered jellyfish with biomimetic propulsion. Nat. Biotech. 30, 792–797 (2012).

21 Mandal, B. B., Grinberg, A., Gil, E. S., Panilaitis, B. & Kaplan, D. L. High-strength silk protein scaffolds for bone repair. Proc. Natl. Acad. Sci. 109, 7699–7704 (2012).

22 Trachsel, L., Johnbosco, C., Lang, T., Benetti, E. M. & Zenobi-Wong, M. Double-Network Hydrogels Including Enzymatically Crosslinked Poly-(2-alkyl-2-oxazoline)s for 3D Bioprinting of Cartilage-Engineering Constructs. Biomacromolecules 20, 4502–4511 (2019).

23 Butcher, A. L., Offeddu, G. S. & Oyen, M. L. Nanofibrous hydrogel composites as mechanically robust tissue engineering scaffolds. Trends Biotech. 32, 564–570 (2014).

24 Cr Keese, I. G. Cell Growth on Liquid Microcarriers. Science 219, 1448–1449 (1983).

25 Giaever, I. & Keese, C. R. Behavior of cells at fluid interfaces. Proc. Natl. Acad. Sci. 80, 219–222 (1983).

26 Kong, D. et al. Protein nanosheet mechanics controls cell adhesion and expansion on low-viscosity liquids. Nano Lett. 18, 1946–1951 (2018).

27 Kong, D., Nguyen, K. D. Q., Megone, W., Peng, L. & Gautrot, J. E. The culture of HaCaT cells on liquid substrates is mediated by a mechanically strong liquid-liquid interface. Faraday Discuss. 204, 367–381 (2017).

28 Minami, K. et al. Suppression of myogenic differentiation of mammalian cells caused by fluidity of a liquid-liquid interface. Appl. Mater. Interfaces 9, 30553–30560 (2017).

29 Peng, L. & Gautrot, J. Long term expansion profile of mesenchymal stromal cells at protein nanosheet-stabilised bioemulsions for next generation cell culture microcarriers. Mater. Today Bio 12, 100159 (2021).

30 Kong, D., Peng, L., di Cio, S., Novak, P. & Gautrot, J. E. Stem cell expansion and fate decision on liquid substrates are regulated by self-assembled nanosheets. ACS Nano 12, 9206–9213 (2018).

31 Jia, X. et al. Adaptive Liquid Interfacially Assembled Protein Nanosheets for Guiding Mesenchymal Stem Cell Fate. Adv. Mater., 1905942 (2019).

32 Bos, M. A. & van Vliet, T. Interfacial rheological properties of adsorbed protein layers and surfactants: a review. Adv. Colloid Interf. Sci. 91, 437–471 (2001).

33 Fuller, G. G. & Vermant, J. Complex fluid-fluid interfaces: rheology and structure. Annu. Rev. Chem. Biomol. Eng. 3, 519–543 (2012).

34 Bergfreund, J. et al. Globular protein assembly and network formation at fluid interfaces: effect of oil. Soft Matter 17, 1692–1700 (2021).

35 Felix, M., Romero, A., Sanchez, C. C. & Guerrero, A. Modelling the non-linear interfacial shear rheology behavioiur of chickpea protein-adsorbed complex oil/water layers. Appl. Surf. Sci. 469, 792–803 (2019).

36 Chang, C.-H. & Franses, E. I. Adsorption dynamics of surfactants at the air/water interface: a critical review of mathematical models, data, and mechanisms. Coll. Surf. A 100, 1–45 (1995).

37 Kong, D. et al. Impact of the Multiscale Viscoelasticity of Quasi-2D Self-Assembled Protein Networks on Stem Cell Expansion at Liquid Interfaces. Biomaterials in Press (2022).

38 Pace, C. J. & Gao, J. Exploring and Exploiting Polar π Interactions with Fluorinated Aromatic Amino Acids. Acc. Chem. Res. 46, 907–915 (2013).

39 Iza, M. & Bousmina, M. Damping function for narrow and large molecular weight polymers: comparison with the force-balanced network model. Rheol. Acta 44, 372–378 (2005).

40 Doi, M. & Edwards, S. F. The theory of polymer dynamics. (1986).

41 Soskey, P. R. & H, W. H. Large step shear strain experiments with parallel-disk rotational rheometers. J Rheol. 28, 625–645 (1984).

42 Tan, K. Y., Gautrot, J. E. & Huck, W. T. S. Formation of pickering emulsions using ion-specific responsive colloids. Langmuir 27, 1251–1259 (2011).

43 Krishnamoorthy, M. et al. Solution conformation of polymer brushes determines their interactions with DNA and transfection efficiency. Biomacromolecules 18, 4121–4132 (2017).

44 Qu, F., Li, D., Ma, X., Chen, F. & Gautrot, J. E. A kinetic model of oligonucleotide-brush interactions for the rational design of gene delivery vectors. Biomacromolecules 20, 2218–2229 (2019).

45 Rodriguez-Maldonado, L., Fernandez-Nieves, A. & Fernandez-Barbero, A. Dynamic light scattering from high molecular weight poly-L-lysine molecules. Coll. Surf. A 270-271, 335–339 (2005).

46 Adamczyk, Z., Morga, M., Kosior, D. & Batys, P. Conformations of Poly-L-lysine Molecules in Electrolyte Solutions: Modeling and Experimental Measurements. J. Phys. Chem. C 122, 23180–23190 (2018).

47 Creton, C., Brown, H. R. & Shull, K. R. Molecular weight effects in chain pullout. Macromolecules 27, 3174–3183 (1994).

48 Creton, C., Kramer, E. J., Hui, C.-Y. & Brown, H. R. Failure mechanisms of polymer intefaces reinforced with block copolymers. Macromolecules 25, 3075–3088 (1992).

49 Ritchie, R. O. The conflicts between strength and toughness. Nat. Mater. 10, 817–822 (2011).

50 Zhao, X. Multi-scale multi-mechanism design of tough hydrogels: building dissipation into stretchy networks. Soft Matter 10, 672–687 (2014).

51 Gong, J. P. Why are double network hydrogels so tough? Soft Matter 6, 2583–2590 (2010).

52 Discher, D. E., Janmey, P. & Wang, Y.-L. Tissue cells feel and respond to the stiffness of their substrate. Science 310, 1139–1143 (2005).

53 Trappmann, B. et al. Extracellular matrix tethering regulates stem cell fate. Nat. Mater. 11, 642–649 (2012).

54 Trichet, L. et al. Evidence of a large-scale mechanosensing mechanism for cellular adaptation to substrate stiffness. Proc. Natl. Acad. Sci. 109, 6933–6938 (2012).

55 Tan, J. L. et al. Cells lying on a bed of microneedles: an approach to isolate mechanical force. Proc. Natl. Acad. Sci. 100, 1484–1489 (2003).

56 Gautrot, J. E. et al. The nanoscale geometrical maturation of focal adhesions controls stem cell differentiation and mechanotransduction. Nano Lett. 14, 3945–3952, doi:10.1021/nl501248y (2014).

57 Di Cio, S., Iskratsch, T., Connelly, J. T. & Gautrot, J. E. Contractile myosin rings and cofilin-mediated actin disassembly orchestrate ECM nanotopography sensing. Biomaterials 232, 119683 (2020).

58 Carisey, A. et al. Vinculin regulates the recruitment and release of core focal adhesion proteins in a force-dependent manner. Curr. Biol. 23, 273–281 (2013).

