## Supplementary Figures and Methods for "Mesenchymal Stem Cells Sense the Toughness of Nanomaterials and Interfaces"

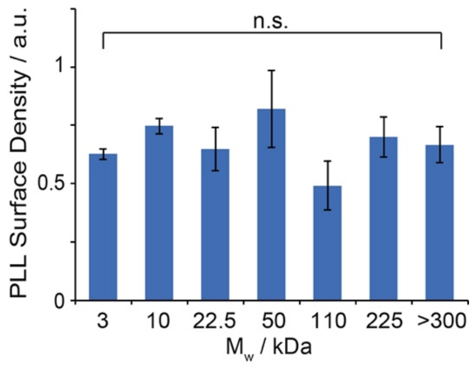


**Supplementary Figure S1.** Quantification of PLL density within nanosheets assembled at the Novec 7500-water interfaces (Novec 7500 containing 10 µg/mL PFBC; aqueous solution is PBS with pH adjusted to 10.5). PLL (tagged with fluorescent dye) with different $M_w$ (3, 10, 22.5, 50, 110, 225 and >300 kDa) was introduced to make a final solution with a concentration of 100 µg/mL.

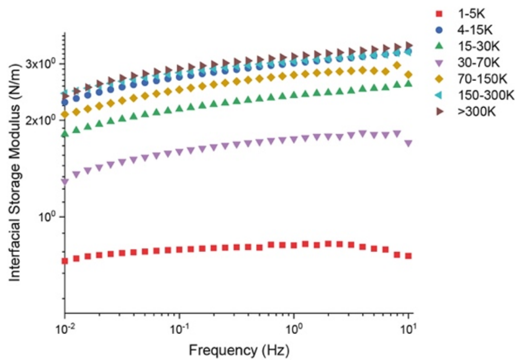


**Supplementary Figure S2.** Frequency sweep profiles of interfaces consisting of PLL nanosheets assembled at the Novec 7500-water interfaces (Novec 7500 containing 10 µg/mL PFBC; aqueous solution is PBS with pH adjusted to 10.5; strain of $10^{-3}$ rad). PLL with different $M_w$ (3, 10, 22.5, 50, 110, 225 and >300 kDa) was introduced to make a final solution with a concentration of 100 µg/mL.

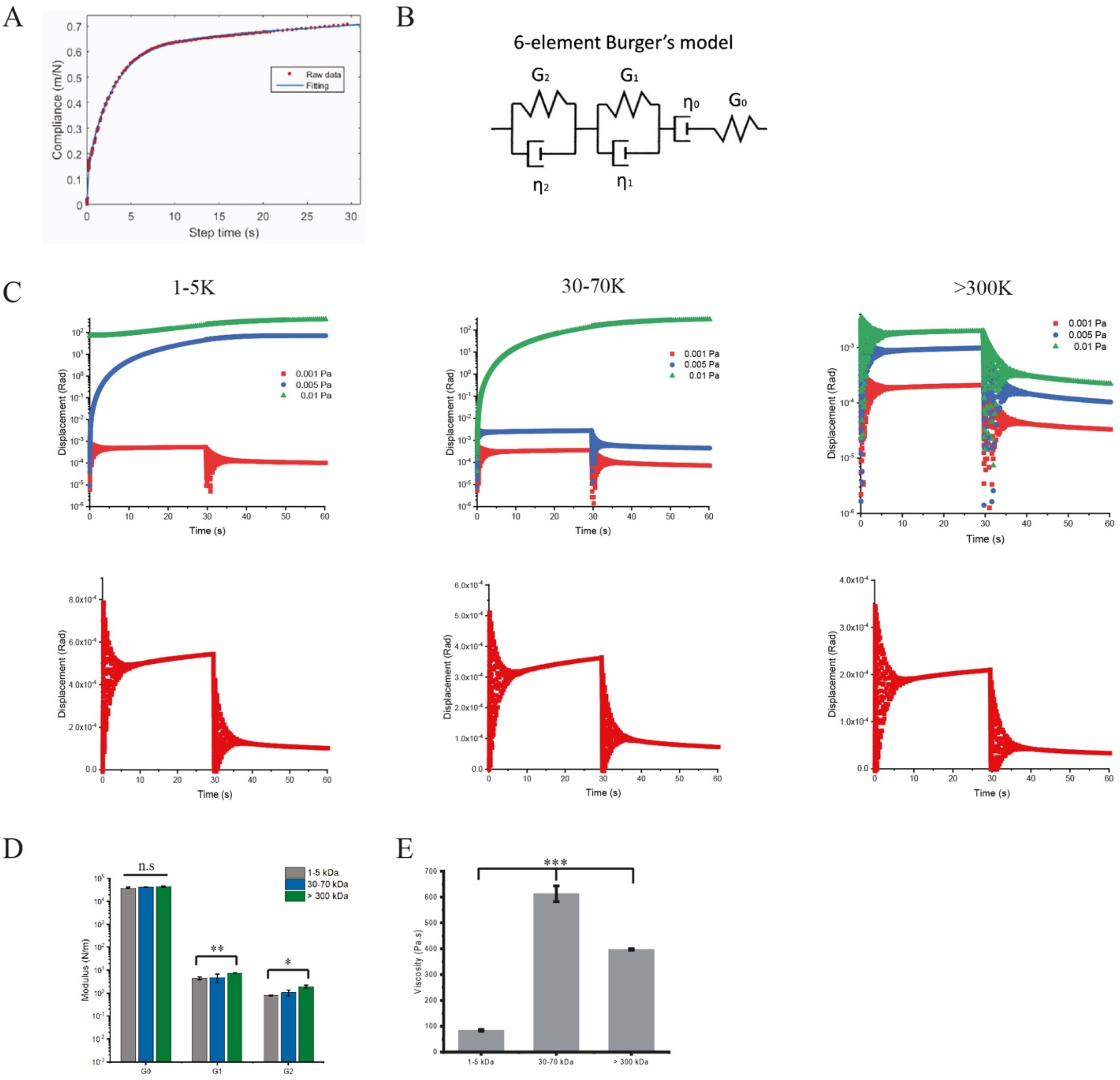


**Supplementary Figure S3.** Interfacial creep experiments carried out at the Novec 7500/PBS interfaces (Novec supplemented with 10 μg/mL PFBC; PBS with pH adjusted to 10.5, containing 100 μg/mL PLL at different $M_w$ (3, 50 and >300 kDa). A-B) Representative creep trace fitted with a 6-element Burger's model and schematic illustration of a 6-element Burger's model. It consists of two Kelvin–Voigt elements (a spring and a dashpot in parallel) in series with a Maxwell element (a dashpot and a spring in series). C) Representative creep and recovery curves at different oscillation stresses (top) and at stress of 0.001 Pa alone (bottom). D-E) The viscoelasticity parameters (D, elastic modulus, G0, G1 and G2, top; E, and viscosity η) extracted from interfacial creep experiments, fitted with 6-element Burger's model. Error bars are s.e.m.; n=3.

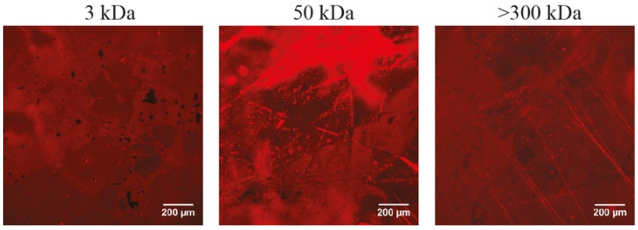


**Supplementary Figure S4.** Epifluorescence images of PLL nanosheets harvested from liquid-liquid interfaces on mica substrates, via Langmuir Blodgett deposition and confirming that large areas are fully covered by nanosheets.

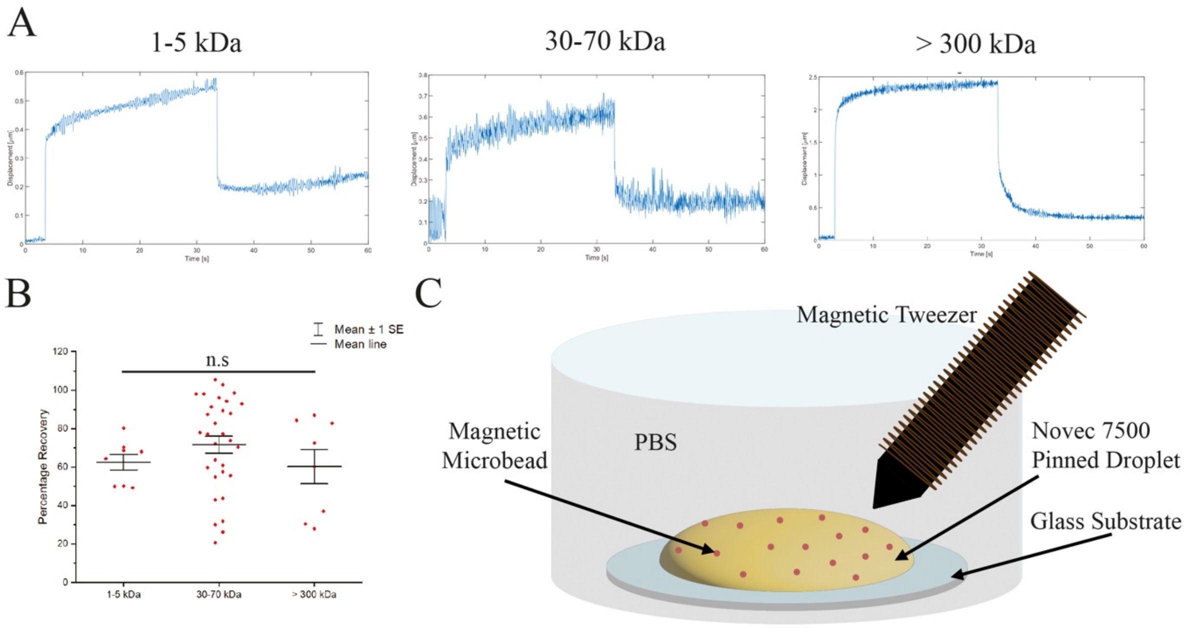


**Supplementary Figure S5.** Interfacial microrheology creep experiments using magnetic tweezers. Magnetic beads were allowed to adhere to nanosheets formed at the interface between Novec 7500 containing 10 µg/mL PFBC and PBS (pH 10.5) containing 100 µg/mL PLL at different $M_w$ (3, 50 and >300 kDa). (A) Representative creep and recovery curves and (B) corresponding percentage recovery extracted from fitting the data to a 6-element Burger's model. Error bars are s.e.m.; n≥7. (C) Schematic representation of the magnetic tweezer experimental set up for magnetic tweezer in this study.

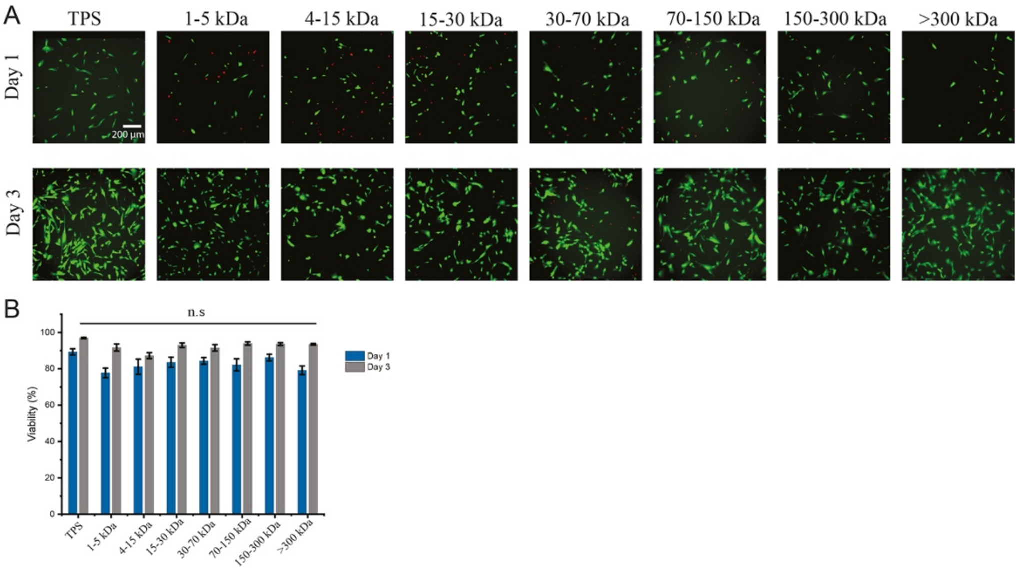


**Supplementary Figure S6.** A) Epifluorescence microscopy images of MSCs cultured on TPS and Novec 7500 oil interfaces, stabilised by PLL nanosheets (days 1 and 3). Live/Dead staining: green, live cells; red, dead cells. B) Quantification of corresponding cell viabilities. Error bars are s.e.m,; n $\geqslant$ 3. Detail of interfaces: Novec 7500 containing 10 μg/mL PFBC; aqueous solution is PBS with pH adjusted to 10.5; PLL with different $M_w$ (3, 10, 22.5, 50, 110, 225 and >300 kDa) at a final concentration of 100 μg/mL.

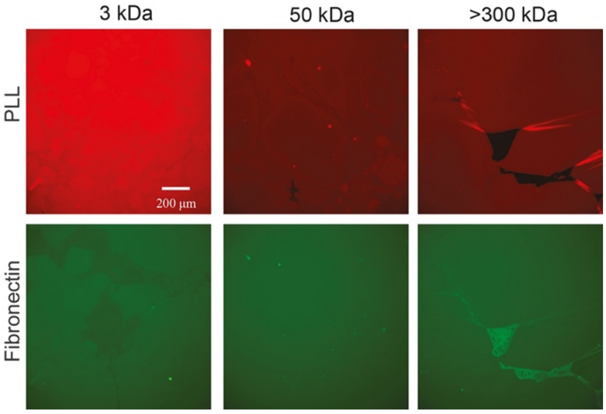


**Supplementary Figure S7.** Epifluorescence images of PLL and fibronectin forming PLL/fibronectin nanosheets assembled at the interfaces between Novec 7500 containing 10 μg/mL PFBC and PBS (pH adjusted to 10.5; PLL with different $M_w$ (3, 10, 22.5, 50, 110, 225 and >300 kDa) at a final concentration of 100 μg/mL.

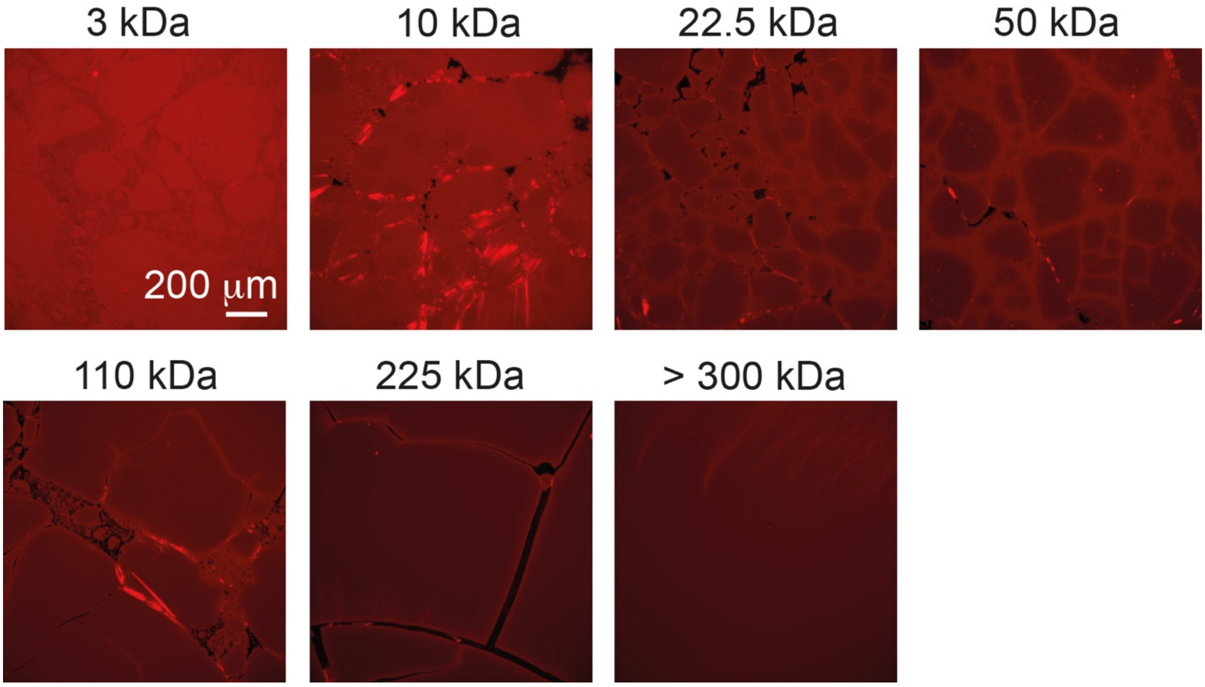


**Supplementary Figure S8.** Highly confluent MSCs remodel and fracture PLL/FN nanosheets assembled at the surface of Novec 7500. Epifluorescence microscopy images of PLL nanosheets after fibronectin coating, prior to cell seeding. Detail of interfaces: Novec 7500 containing 10 μg/mL PFBC; aqueous solution is PBS with pH adjusted to 10.5; PLL with different $M_w$ (3, 22.5, 110 and 225 kDa) at a final concentration of 100 μg/mL.

### Supplementary Materials and Methods

**Interfacial rheology.** Interfacial rheology was carried out on a hybrid rheometer (DHR-3) from TA Instruments fitted with a double wall ring (DWR) geometry and a Delrin trough with a circular channel. The double wall ring used for this geometry has a radius of 34.5 mm and the thickness of the Platinum–Iridium wire is 1 mm. The diamond-shaped cross-section of the geometry's ring provides the capability to pin directly onto the interface between two liquids and measure the interface properties without sub-phase correction. 19 mL of the fluorinated oil (Novec 7500, ACOTA) pre-mixed with pentafluorobenzoyl chloride (PFBCl, Sigma-Aldrich) at desired concentrations was placed in the Delrin trough and the ring was lowered, ensuring contact with the surface, via an axial force procedure. The measuring position was set 500 µm lower than the contact point of the ring with the oil-phase surface. Thereafter, 15 mL of the pH 10.5 PBS buffer was carefully syringed on top of the oil phase. Time sweeps were performed at a constant frequency of 0.1 Hz and a temperature of 25 °C, with a displacement of 1.0 $10^{-3}$ rad to follow the formation of the protein layers at the interface. The concentration of poly(L-lysine) (PLL) used for all rheology experiments were 100 µg/mL (with respect to aqueous phase volume), respectively. Before and after each time sweep, frequency sweeps (with a constant displacement of $10^{-3}$ rad) were conducted to examine the frequency-dependant characteristics of the interface.

Before amplitude sweeps (with constant frequencies of 0.1 Hz) were carried out, stress relaxation was performed at 1% strain for 120 s. Considering the low moduli initially measured for pristine liquid-liquid interfaces (in the absence of protein and/or surfactant), viscous drag from both phases were not corrected. We note that although the interfacial shear moduli observed at liquid-liquid interfaces in the absence of protein or surfactant are expected to be considerably lower than those measured in our assay (*1, 2*), due to lack of viscous drag correction, they should not completely vanish and interfacial shear viscosity should be expected to persist, even with liquid-liquid interfaces that display very limited roughness (at the molecular scale). Strain sweeps (constant frequencies of 0.1 Hz) were caried out last, as typically resulting in fracture/disruption of interfaces at end points. Damping functions were generated by normalising strain sweep data to moduli measured at the lowest strain. To quantify interfacial toughness, corresponding stress-strain plots were integrated. We note that these interfacial toughness data, extracted from such shear experiments, cannot be quantitatively compared to toughness data extracted from tensile testing, even after integration of interfacial toughness to the thickness of nanosheets.

**Data analysis from interfacial stress relaxation experiments.** Stress relaxation data from the 10$^{th}$ second onwards when the stress relaxation started was plotted as stress against time in scatter in OriginPro, and fitted with double exponential decay fit, according to the following equation:

$$\sigma = \sigma_e + \sigma_1\left(1 - e^{-t/\tau_1}\right) + \sigma_2\left(1 - e^{-t/\tau_2}\right) \qquad (1)$$

In this equation, $\sigma$ is the measured residual stress, $\sigma_e$ is the elastic stress and $\sigma_1$ and $\sigma_2$ are viscous relaxation components. The degree of stress retention ($\sigma_r$) is calculated as:

$$\sigma_r = \frac{\sigma_e}{\sigma_e+\sigma_1+\sigma_2} \times 100 \qquad (2)$$

**Interfacial creep experiments.**

For creep recovery experiments, a double wall Du Noüy Ring geometry (20 mm in diameter and 400 µm in thickness) and trough of corresponding dimensions were used. 3 mL of the fluorinated oil pre mixed with 10 µg/mL PFBC were placed in the Delrin trough and the ring was lowered, ensuring contact with the surface, via an axial force procedure. The measuring position was set 200 µm lower than the contact point of the ring with the oil phase surface. Thereafter, 4 mL of the pH 10.5 PBS buffer was carefully syringed on top of the oil phase. Time sweep was performed at a constant frequency of 0.1 Hz and a temperature of 25°C, with a displacement of $10^{-3}$ rad to follow the formation of the protein layers at the interface. A 40 µL amount of 10 mg/mL (in DI water) PLL with molecular weight of interest was pipetted to the aqueous phase, making a final concentrations at 100 µg/mL. When the time sweep measurement was completed, the excess polymer solution was washed by diluting with PBS (pH 7.4) six times. The creep recovery experiment was then carried out by applying a $10^{-3}$, 5 $10^{-3}$ or $10^{-4}$ Pa stress for a duration of 30 s, followed by a 30 s recovery at 0 stress. Due to creep ringing artefacts resulting from the coupling of the instrument inertia and sample elasticity in a stress controlled rheometer (*3-5*), smoothing of traces was applied in Origin and the creep results were fitted with a 6 element modified Burger's model in MATLAB, according to the following equation:

$$J = \frac{1}{G_0} + \frac{1}{G_1}\left(1 - e^{-t/\tau_1}\right) + \frac{1}{G_2}\left(1 - e^{-t/\tau_2}\right) + \frac{t}{\eta} \qquad (3)$$

Where *J* is the creep compliance, whereas $G_0$, $G_1$, $G_2$, are the elastic and viscous shear moduli (two relaxation time components), and $\tau_1$, $\tau_2$ and $\eta$ are the relaxation times and viscosity of the corresponding interfaces.

**Generation of pinned droplets.** Glass samples were placed into a desiccator together with an open vial containing toluene (1 mL) and 30 µL trichloro (1H, 1H, 2H, 2H-perfluorooctyl) silane (Sigma). The desiccator was placed under vacuum for 5 min and then left under reduced atmosphere but sealed overnight. After 24 h incubation, the glass slides were washed with ethanol and dried in air. The resulting hydrophobic glass slides were cut into 1×1 cm samples and placed into a 24-well plate. After sterilization with 70% ethanol, samples were washed with PBS and filled with 1 mL PBS (with pH adjusted as indicated). 100 µL pinned droplets of fluorinated oil (Novec 7500) with the fluorinated co-surfactant at desired concentrations were deposited on top of the submerged coated glass slide. 1 mL of 200 µg/mL PLL solution was pipetted into each well (final concentration of 100 µg/mL) and left to incubate for 1 h. Samples were washed by successive dilution/aspiration with PBS (pH 7.4, 6 times). Fibronectin adsorption was carried out by adding 20 µL of a fibronectin solution (1 mg/mL) into each well (final concentration: 10 µg/mL), followed by incubation at room temperature for 1 h. Finally, samples were washed by dilution/aspiration with PBS (pH 7.4) four times and then with growth medium twice.

**Interfacial creep microrheology.** *Preparation of samples.* Fluorinated glass slides (prepared as above) were placed into a 3 cm petri-dish and covered with 5 mL of PBS (normally at pH 10.5 unless otherwise specified). Pinned droplets of 100 µL fluorinated oil with PFBCl at desired concentrations were deposited at the fluorinated surface, covering the entire substrate. 50 µL PLL solution (10 mg/mL) was pipetted into the dish, making a final concentration of 100 µg/mL. After 1 h incubation, the polymer solution was washed by sequential dilution/aspiration with PBS (pH 7.4) six times. Meanwhile, epoxylated dynabeads (4.5 µm, Thermo Fisher) were diluted in PBS (100 times from stock), and 50 µL of the resulting suspension was homogeneously pipetted onto the dish and left to incubate for 15 min. The excess beads in the dish were removed by sequential dilution/aspiration with PBS (pH 7.4) six times, priot to the start of measurements.

*Magnetic tweezer-operated interfacial creep microrheology.* Interfacial creep-recovery experiments were performed using magnetic tweezers. Beads bound to the nanosheets were subjected to a force pulse with a magnitude of 6 nN (40 µm distance between the bead and the tip) and a duration of 30 s. Their trajectories were recorded for a total of 1 minute per bead with a frame every 20 ms. Around 30 traces were obtained per dish, per experiment. Bead tracking was achieved by MATLAB and fitted with a 6-element modified Burger's model (two Kelvin–Voigt elements in series with a Maxwell element, extended from a 4-element model (*6, 7*)), according equation (3) and defining parameters as follows:

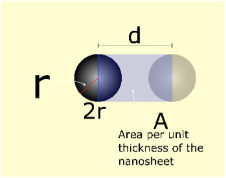


$$\text{Strain} = \frac{d}{r}$$

$$\text{Stress} = \frac{F}{A} = \frac{F}{2r}$$

$$J = \frac{\text{Strain}}{\text{Stress}} = \frac{2d}{F}$$

For $G_0$, n = 11, 8, 7 for low, medium and high $M_w$, respectively; for $G_1$, n = 10, 8, 7 for low, medium and high $M_w$, respectively; for $G_2$, n = 9, 8, 8 for low, medium and high $M_w$, respectively.

**X-ray photoelectron spectroscopy.** Emulsions generated between Novec 7500/PFBCl (10 µg/mL) and PLL aqueous solutions (in PBS, pH 10.5, at a concentration of 100 µg/mL; 1/2 fluorinated oil to aqueous solution ratio) were washed 9 times with deionised water and allowed to dry on silicon substrates. Dried protein nanosheets were washed with hexafluoroisopropanol (Sigma) and ethanol to remove any soluble residues, and finally dried again. XPS was carried out using a Nexsa X-ray Photoelectron Spectrometer (XPS) System on samples prepared as for SEM characterisation. A pass energy of 200 eV and a step size of 1 eV were used for survey spectra. For high energy resolution spectra, a pass energy of 50 eV and a step size of 0.1 eV were used. The spectrometer charge neutralising system was used to compensate sample charging and the binding scale was referenced to the aliphatic component of C 1s spectra at 285.0 eV. The concentrations obtained are reported as the average percentage of that particular atom species (atomic %) at the surface of 4 samples (< 10 nm analysis depth) without any correction. The analysis area ($0.3 \times 0.7$ mm$^2$), the angle of incidence and the beam intensity were kept constant for all measurements.

**Mesenchymal stem cells (MSCs) culture and seeding.** Bone marrow derived human mesenchymal stem cells (PromoCell) were cultured on T75 flasks in MSC growth medium (PromoCell). MSCs were harvested with 4 mL accutase-solution (PromoCell), resuspended, then centrifuged. 5,000 cells per well (resuspended in medium) were seeded on flat interfaces (per well in 24 well plates) and cultured in an incubator (37°C and 5% $CO_2$). Half of the medium was replaced with fresh medium every two days. For passaging, 300,000 cells were seeded in a T75 flask.

**Generation of flat fluorinated oil-culture medium interfaces for monitoring of cell expansion.** 24 well-plates were plasma treated for 10 min. 500 µL of ethanol containing 10 µL trimethylamine and 10 µL trichloro (1H, 1H, 2H, 2H-perfluorooctyl) silane were added into each well. After 24 h incubation, the solution was removed and wells were washed with ethanol. After washing with PBS twice, 500 µL of Novec 7500 containing the desired prosurfactant at a desired concentrations (see detail of each figure) were transferred into each well. 1 mL of PBS (pH adjusted to 10.5 ) were carefully pipetted at the surface of the oil, followed by 1 mL of 200 µg/mL PLL solution (in pH 10.5 PBS; final concentration of 100 µg/mL) and incubation for 1 h. Interfaces were washed by dilution/aspiration with PBS (pH 7.4) six times. Fibronectin adsorption was carried out by adding 20 µL of a fibronectin solution (1 mg/mL) into each well, to make a final concentration of 10 µg/mL, and incubated for 1 h. Functionalised interfaces were washed by dilution/aspiration with PBS (pH 7.4) four times, and then with growth medium twice.

**Preparation of glass-mounted wells for higher resolution imaging.** Fluorinated thin glass slide (25×60 mm) was attached to Sticky-Slide 8 Well (An 8 well bottomless µ-Slide with a self-adhesive underside to which substrates can be mounted, Ibidi). After sterilization with 70% ethanol, wells were washed with PBS and filled with 600 µL PBS (with pH adjusted as indicated). Then 10 µL pinned droplets of fluorinated oil (Novec 7500) with the fluorinated co-surfactant (PFBCl) at desired concentrations was deposited on top of the submerged coated glass substrate. 300 µL PBS was removed by micropipette aspiration. 300 µL of 200 µg/mL PLL solution was pipetted into each well (final concentration of 100 µg/mL) and left to incubate for 1 h. Each well was washed by successive dilution/aspiration with PBS (pH 7.4, 6 times). Fibronectin adsorption was carried out by adding 6 µL of a fibronectin solution (1 mg/mL) into each well (final concentration: 10 µg/mL), followed by incubation at room temperature for 1 h. Finally, each well was washed by dilution/aspiration with PBS (pH 7.4) four times and then with growth medium twice.

**Hoechst staining and cell counting.** Cell proliferation on flat interfaces was assessed via Hoechst staining. Half of the medium was replaced by pre-warmed PBS containing 2 µL Hoechst (1 mg/mL Thermofisher Scientific). After 30 min incubation, cells were imaged using a Leica DMI4000 fluorescence or or Leica DMi8 epifluorescence microscope. Cell counting was carried out by thresholding and watershedding nuclear images in ImageJ.

**Immunostaining.** For immunostaining and imaging at higher resolution, cells were cultured on pinned droplet generated in a sticky-slide 8 well plates (Ibidi), prepared as described above. After 24 or 48 h incubation, each well was diluted with PBS six times before samples were fixed with 8 % paraformaldehyde for 10 min and diluted with PBS six times before permeabilization with 0.4% Triton X-100 for 5 min at room temperature. Samples were blocked for 1 h (blocking buffer: PBS containing 10 vol% foetal bovine serum and 0.5 vol% gelatine), combining with tetramethyl rhodamine isothiocyanate phalloidin (1:500, Sigma-Aldrich). Samples were subsequently incubated with primary antibodies (anti-vinculin mouse monoclonal, Sigma-Aldrich, 1:200 in blocking buffer) for 1 h at room temperature, diluted with PBS six times, then incubated with Alexa Fluor 488-conjugated secondary antibodies (goat anti-mouse, 1:500 in blocking buffer) and DAPI (1:500) for 1 h at room temperature. The samples were washed six times by dilution with deionised water and imaged shortly after.

**Imaging of PLL nanosheets assembled at oil interfaces.** Fluorinated glass slides (1 x 1 cm) were placed in a 24 well plate. After sterilisation with 70 % ethanol, the wells were washed (twice) and then filled with 2 mL PBS (pH 10.5). 100 µL droplets of fluorinated oil (Novec 7500) with the fluorinated co-surfactant PFBCl at a concentration of 10 µg/mL were deposited on the glass samples and formed a fluorinated oil droplet spreading over the entire substrate. Subsequently, a labelled PLL solution (2 µL,

PLL-Alexa Fluor$^{TM}$ 594 at 10 mg/mL, mixed with 18 µL of PLL solution at 10 mg/mL) was added to PBS to make a final PLL concentration of 100 µg/mL, and the resulting interfaces were incubated for 30 min. PLL adsorption was interrupted by reducing the pH below 5, by adding a drop of 1.0 M HCl. The staining solution was then diluted with PBS (pH 7.4) 8 times, prior to fluorescence imaging.

**Immuno-fluorescence microscopy and data analysis.** Fluorescence microscopy images were acquired with a Leica DMI4000B fluorescence microscopy (CTR4000 lamp; 63 × 1.25 NA, oil lens; 10 × 0.3 NA lens; 2.5 x 0.07 NA lens; DFC300FX camera) and a Leica DMi8 epifluorescence microscope (HC PL FLUOTAR 10x/0.32 PH1; HC PL FLUOTAR 63X/1.30 Oil PH3; LEICA DFC9000 GT sCMOS camera). Confocal microscopy images were acquired with a Leica TCS SP2 confocal microscope (X-Cite 120 LED lamp; 63 x 1.40-0.60 NA, oil lens; 10 x 0.3 NA lens; DFC420C CCD camera) and a Zeiss Super resolution LSM710 ELYRA PS.1 (EC Plan-Neofluar10x/0.3 M27; EC Plan-Neofluar20x/0.5 M27; sCMOS camera). Cell densities were determined after thresholding and watershedding nuclei images in ImageJ. In the case of cell aggregates, cells were counted manually. To determine adhesion cell areas, images were analysed by outlining the contour of the cell cytoskeleton (phalloidin stained) and areas were measured in ImageJ. To determine cell spreading areas, images were analyzed by thresholding and watershedding cytoskeleton images (phalloidin staining). For confocal imaging, stacks of 16 sections were scanned, with an image averaging of 2 and a line averaging of 4. 3D reconstruction and volume rendering of the stacks were performed via Imaris x64.

**Statistical analysis.** Statistical analysis was carried out using Origin 2019 through one-way ANOVA with Tukey test for posthoc analysis. Significance was determined by * $P < 0.05$, ** $P < 0.01$, *** $P < 0.001$ and n.s., non-significant. A full summary of statistical analysis is provided below as a separate supporting file.


1. Z. A. Zell *et al.*, Surface shear inviscidity of soluble surfactants. *Proc. Natl. Acad. Sci.* **111**, 3677-3682 (2014).
2. S. Vanderbril, A. Franck, G. G. Fuller, P. Moldenaers, J. Vermant, A double wall-ring geometry for interfacial shear rheometry. *Rheol. Acta* **49**, 131-144 (2010).
3. T. B. Goudoulas, N. German, Viscoelastic properties of polyacrylamid solutions from creep ringing data. *J. Rheol.* **60**, 491-502 (2016).
4. L. Pavlovsky, J. G. Younger, M. J. Solomon, In situ rheology of staphylococcus epidermidis bacterial biofilms. *Soft Matter* **9**, 122-131 (2013).
5. A. Jaishankar, V. Sharma, G. H. McKinley, Interfacial viscoelasticity, yielding and creep ringing of globular protein-surfactant mixtures. *Soft Matter* **7**, 7623-7634 (2011).
6. O. Chaudhuri *et al.*, Substrate stress relaxation regulates cell spreading. *Nat. Commun.* **6**, 6365 (2015).
7. M. Dogan, A. Kayacier, O. S. Toker, M. T. Yilmaz, S. Karaman, Steady, Dynamic, Creep, and Recovery Analysis of Ice Cream Mixes Added with Different Concentrations of Xanthan Gum. *Food Bioprocess. Technol.* **6**, 1420-1433 (2013).
